# Head MagnetoMyography (hMMG): a novel approach to monitor face and whole-head muscular activity

**DOI:** 10.1101/556100

**Authors:** Guido Barchiesi, Gianpaolo Demarchi, Frank H. Wilhelm, Anne Hauswald, Gaëtan Sanchez, Nathan Weisz, Guido Barchiesi

## Abstract

Muscular activity recording is of high basic science and clinical relevance and is typically achieved using electromyography (EMG). While providing detailed information about the state of a specific muscle, this technique has limitations such as the need for a-priori assumptions about electrode placement and difficulty with recording muscular activity patterns from extended body areas at once. For head and face muscle activity, the present work aimed to overcome these restrictions by exploiting magnetoencephalography (MEG) as a whole-head myographic recorder (head magnetomyography, hMMG). This is in contrast to common MEG studies, which treat muscular activity as artifact in electromagnetic brain activity. In a first proof-of-concept step, participants imitated emotional facial expressions performed by a model. Exploiting source projection algorithms, we were able to reconstruct muscular activity, showing spatial activation patterns in accord with the hypothesized muscular contractions. Going one step further, participants passively observed affective pictures with negative, neutral, or positive valence. Applying multivariate pattern analysis to the reconstructed hMMG signal, we were able to decode above chance the valence category of the presented pictures. Underlining the potential of hMMG, a searchlight analysis revealed that generally neglected neck muscles exhibit information on stimulus valence. Results confirm the utility of hMMG as a whole-head electromyographic recorder to quantify muscular activation patterns including muscular regions that are typically not recorded with EMG. This key advantage beyond conventional EMG has substantial scientific and clinical potential.

## 1 Introduction

Our body allows for expression of a rich set of emotional states intrinsic to social interaction (Darwin, 1872; Nummenmaa, Glerean, Hari, & Hietanen, 2014; Nummenmaa, Hari, Hietanen, & Glerean, 2018). Especially facial muscles have a fundamental role in communicating emotion and mood, i.e., expressing and, from the observer point of view, perceiving, related bodily signals. To investigate emotional expression, studies in this field often exploit a technique of muscular recording called surface electromyography (EMG; Fridlund & Cacioppo, 1986; Larsen, Norris, & Cacioppo, 2003). When expressing a specific emotional state, specialized muscles in the face contract, resulting in differentiated activity patterns of facial muscles. In order to start a muscle contraction, motor neurons release acetylcholine at the neuromuscular junction leading to muscle fiber action potential, which in turn causes the fiber to contract. Typically EMG uses a bipolar arrangement to record the voltage difference between two electrodes placed along the muscle fibers direction. Given its low cost and relative ease of use, EMG is the gold standard for non-invasive recordings of muscular electrical activity. While yielding signals at relatively high signal-to-noise levels for a specific muscle, a limitation is that, in order to record different muscles, many electrodes are needed. Naturally spatial and practical limits are reached fairly quickly (commonly not more than 5-6 muscles are investigated) forcing the researchers to an a-priori selection. Further, some muscles are very difficult to record via surface EMG (e.g., inner neck). These issues limit the clinical use of the surface and (more invasive) needle EMG, since indeed some muscles (such as *Longus Colli* in chronic neck pain syndromes, Falla, Jull, O’Leary, & Dall’Alba, 2006) are difficult to record with these techniques. In addition, the use of conventional techniques can be highly unpleasant in pain conditions such as allodynia or hyperalgesia (Coutaux, Adam, Willer, & Le Bars, 2005).

In an attempt to overcome these issues, we introduce a novel application of whole-head MEG, to record and reconstruct myographic activity from the head at once. MEG uses superconductive sensors (SQUIDs) to record magnetic potentials primarily produced by postsynaptic currents. Given the resemblance between functioning principles of neurons and muscle fiber contraction, MEG also records electromagnetic signals originating from muscles. In fact, in typical cognitive neuroscience experiments, muscular activity is visible even in MEG raw signals. Such muscular signals are normally regarded as noise that needs to be removed from the neural data. In the present work we treat muscular activity as signal, since MEG has distinct advantages with respect to the aforementioned EMG-related issues: a) no electrodes and no a-priori locations are needed since MEG records from the entire head at once; b) ideally, it might also record deep muscle activity, which (using inverse models) could be separated from more superficial ones.

Already fifty years ago researchers attempted to record magnetic fields generated from muscular contractions: David Cohen was the first, using one SQUID, to record activity from forearm and hand muscles; he called the technique “MagnetoMyography” (MMG, Cohen & Givler, 1972). Nowadays MMG is used to investigate uterine contractions using arrays of SQUIDs specifically implemented for this purposes (Escalona-Vargas, Oliphant, Siegel, & Eswaran, 2019; Eswaran, Preissl, Murphy, Wilson, & Lowery, 2005), and muscular activity from the heart (MagnetoCardiography, Fenici, Brisinda, & Meloni, 2005).

Differently for previous research, the present work focuses on exploiting existing Magnetoencephalography devices to record muscular activity from face and head, and to localize, taking advantage of source reconstruction algorithms, the magnetic activity generated by a variety of face and head muscles at the same time. No one before us, to the best of our knowledge, attempted to apply MMG to localize face and head muscles activity.

In order to make first steps into expanding the application of MMG into the affective neuroscience, we implemented a proof-of-principle that illustrates some of these potential advantages: in a first step, using an imitation task, we demonstrate that the multiple muscular sources contributing to the MEG signal are distinctly localizable on the participants’ face. In a second step, we apply this novel approach to a more typical experimental environment by asking participants to passively watch emotion-eliciting pictures. Validating the approach, we replicate a classical finding in the emotion literature, namely that *m. Corrugator Supercilii* activity correlates negatively with stimulus valence (Cacioppo, Petty, Losch, & Kim, 1986; LANG, GREENWALD, BRADLEY, & HAMM, 1993). Going beyond this replication we use classification algorithms to predict the valence category of the observed picture from the reconstructed muscular signals, pointing to the importance of neck muscles, which has not been considered in studies using conventional EMG. This underscores the promise of this approach as a tool for future scientific discoveries as well as a potential clinical tool. From this point on, we are going to refer to our implementation of MEG muscular recording as head MagnetoMyography (hMMG).

## 2 Methods

### 2.0.1 Participants

The experiment took place in the MEG Lab of the University of Salzburg located at the Christian Doppler Clinics (Salzburg). The protocol was approved by the Ethics Committee of the University of Salzburg. Twenty-two healthy participants (15 females, mean age 24.8 +/- 3.6) were tested, after having signed an informed consent. All participants had normal or corrected-to-normal vision and declared no knowledge of any neurological or psychiatric disorder. Participants received either a monetary reimbursement (10€ per hour), or course credit for their participation. In total the entire experimental session lasted about 2 hours.

### 2.0.2 Data Collection Procedure

Participants were asked to wear a suit and to remove any metal object from their body. Considering the features of the MEG signal we were interested in (gamma to high gamma frequencies, see paragraph 2.1.6), participants wearing metallic braces were not excluded, since the noise produced by braces on the MEG sensors does not affect the frequency bands of interest for the subsequent.

Participants’ head-shape was digitized by means of a Polhemus tracker (3Space FASTRAK, Polhemus, Colchester, US); at least 300 points were tracked on participants’ head, including five head-position (HPI) coils and three anatomical landmarks. For each participant, in addition to the standard head points, also the nose shape and the position of the eyebrows was tracked. On a subset of participants (10) three EMG electrode pairs were placed in correspondence of left *m. Corrugator Supercilii*, right *m. Frontalis*, and right *m. Zygomaticus Major* (Fig. 1A). Participants were then seated in the MEG device in an electromagnetically shielded room (AK3b, Vacuumschmelze, Hanau, Germany). The MEG lab in Salzburg is equipped with a 306-channel Elekta Triux whole-head magnetoencephalography device (102 triplets composed of one magnetometer and two orthogonally placed planar gradiometers). Signal from MEG sensors was sampled at 2000 Hz, with the acquisition filters set to 0.1 Hz high-pass and 660 Hz low-pass. The head position inside the MEG helmet was localized by means of HPI coils, which were energized throughout the whole experiment.

**Fig. 1.**
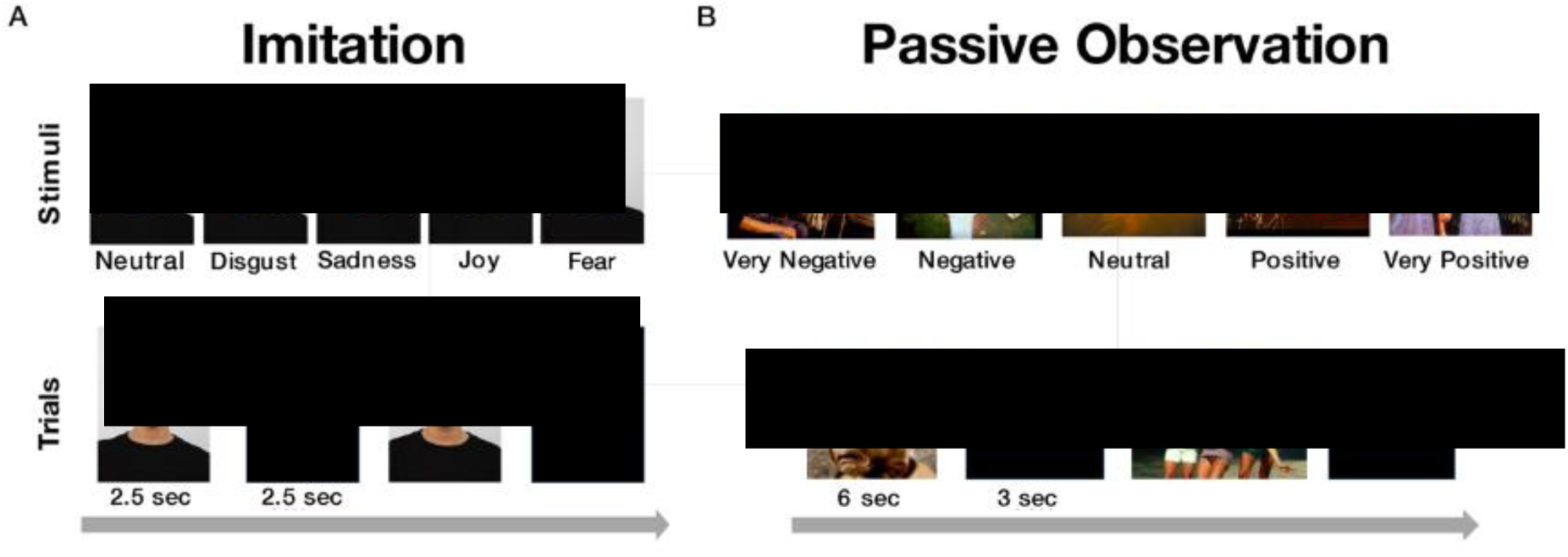
Stimuli and procedure in the Imitation and Passive Observation tasks. **Imitation** (A) *Top*: Pictures showing an actress performing 5 different emotional facial expressions. *Bottom*: Timeline of two trials: pictures were presented for 2.5 seconds, followed by a “relax” label again for 2.5 seconds. (175 pictures, 35 per expression). Participants were asked to imitate the facial expressions observed and to relax their face during the “relax” label. **Passive Observation** (B) *Top*: pictures showing exemplars of the 5 valence categories. *Bottom*: Timeline of two trials: pictures were presented for 6 seconds each, followed by a blank screen for 3 seconds (100 pictures, 20 per valence category). <u>*images have been removed/obscured due to a bioRxiv policy on the inclusion of faces</u>

Visual stimuli were presented by means of a projector (PROPixx, VPixx Saint-Bruno, Canada) back-projecting the images on a semi-transparent screen.

### 2.1 Imitation Task: *Materials and Methods*

#### 2.1.1 Stimuli

We selected 5 pictures from the Radboud Face Database (RaFD, Langner et al., 2010) dataset of an actress producing 5 emotional expressions: Joy, Disgust, Fear, Sadness, Neutral (Fig. 1A, *Top*). Each imitation task session was composed of 175 trials, 35 for each emotion.

#### 2.1.2 Procedure

The presentation of a picture lasted 2.5 seconds on the screen, followed by 2.5 seconds of black screen on which a white “relax” label was presented (Fig. 1A, *Bottom*). The order of the emotion conditions was randomized across participants. Their task was to imitate the emotional expression observed in each trial. Before going inside the MEG scanner, participants were trained outside the shielded room to produce the facial contraction pattern that best approximated the one observed in the pictures. Some participants reported that it was sometimes difficult for them to imitate the facial expressions, especially for fear and sadness pictures. Acquired data were stored on a local server.

#### 2.1.3 EMG Power-Spectrum

EMG power spectrum was computed for the 3 channels recorded from participants wearing EMG electrodes. The signal was band-pass filtered between 9-550 Hz, and then it underwent a time-frequency analysis from 9 to 550 (1 Hz step, 0.3 seconds sliding time-window, “Hanning” taper); the resulting time-frequency data were averaged between 0.5 and 2.5 sec after picture presentation, yielding a single value for each frequency tested. We performed a time-frequency analysis, averaged across time, instead of a classical fast Fourier transform, in order to be consistent with the following analyses. As a final step, trimmed-mean at 20% across trials was computed at each frequency to detect the average power spectrum across participants. A z-score transformation was applied to the power spectra separately for each channel from 9 to 250 Hz. This analysis, besides showing the distribution of power spectra at different muscles, also guided us in selecting a frequency band optimal for the following analyses.

#### 2.1.4 Sensor to Source Space Projection

We conducted two types of analysis for the imitation task, the hMMG-EMG correlation and the hMMG analysis of emotional facial expressions. Both were conducted in source space, projecting the sensor data onto a set of virtual sensors. Fieldtrip toolbox (version updated to November 15, 2018) and custom-made Matlab functions were used to analyze the data (Oostenveld, Fries, Maris, & Schoffelen, 2011). As a first step, we epoched all trials from 0.5 seconds to 2.5 seconds after picture presentation onset. Data were only checked for broken channels, but not for physiologically driven artifacts, given that also participants wearing braces were included in the experiment. Data were band-pass filtered depending on the type of analysis conducted; details on filtering procedures are described in the hMMG-EMG correlation (§ 2.1.5) and the hMMG analysis of emotional facial expressions (§ 2.1.6).

After filtering, sensor data were projected onto source space, separately for each condition, to obtain virtual sensors by multiplying the spatial filters with the single trial data. We obtained structural magnetic resonance images of 10 participants’ heads. For the others we used an MRI model downloaded from the Fieldtrip Toolbox website (ftp:/ftp.fieldtriptoolbox.org/pub/fieldtrip/tutorial/Subject01.zip); we manually (interactively) morphed the MRI shape in order to fit the participant’s head-shape.

Then the morphed head MRIs were segmented into surface (“scalp” tissue) and interior (“brain” tissue). In order to create a volume conduction model, the geometry of the head was determined using a triangulated surface mesh made of 5000 vertices for each compartment. Finally, we used the Fieldtrip implementation of the volume conduction algorithm (“singleshell” option) proposed by Nolte (2003). To create virtual sensors, we scanned the scalp using steps of 1 cm between voxels, and then subtracted the brain volume from it, in order to delete voxels falling into brain tissue. The inner border of the scalp has been extended inside by one centimeter (i.e., setting: cfg.inwardshift = 1), producing a scan grid of 4735 voxels.

Once a volume conduction model and a source-model had been constructed, the leadfield was estimated for each point of the grid and the covariance matrix between MEG sensors was calculated. We applied the *linear constrained minimum variance* (LCMV) beamformer algorithm onto the data (Van Veen, van Drongelen, Yuchtman, & Suzuki, 1997) to generate a spatial filter, mapping virtual sensors to MEG sensors data for each trial. At the end of this procedure we obtained the time-course of the activity of 4735 virtual sensors on the surface of the head.

#### 2.1.5 hMMG-EMG Correlation

Given that projecting data from each single frequency (from 9 to 550 Hz, 1 Hz step, as performed in the EMG Power Spectrum section, § 2.1.3), into source space would have been computationally very demanding, for each participant wearing EMG electrodes we band-pass filtered the data into 9 frequency bands, namely 9-25, 25-45, 45-70, 70-100, 100-160, 160-200, 200-250, 250-350, 450-550 Hz, before projecting them onto source space. To determine which one of the virtual sensors correlates best with each EMG channels, we linearly correlated (Pearson), separately for each participant, for each condition, and for each frequency band, the time-course of each virtual sensor with the time course of each EMG channel, ending up with a matrix of correlation values for each frequency band with number of trials x virtual sensors x number of EMG channels as dimensions (35 x 4735 x 3). We finally averaged the correlation values across trials using a trimmed mean at 20%. Since we were not interested in the direction of the correlation, we computed the absolute value of the obtained matrix.

For each participant and frequency band, absolute correlation values were sorted from the lowest to the highest value. Subsequently, the 2.5% of the virtual sensors showing the highest absolute correlation values were selected. At the group level we computed a map in which every virtual sensor represents the number of participants having that virtual sensor among the 2.5% highest absolute correlation value. Only a subset of condition-EMG pairings were analyzed because of their theoretical correspondence (Cacioppo et al., 1986): Disgust-*Corrugator*, Fear-*Frontalis*, Joy-*Zygomaticus*.

To show the distribution of correlations across frequency bands, we computed a correlation-spectrum for each frequency band. We computed this correlation-spectrum on the voxels showing the highest count and highest count – 1 on the hMMG-EEG correlation maps, separately for each condition (e.g., if maximum count for one condition was 9, then all the channels having 8 or 9 counts were selected). As a final step, separately for each condition, we averaged the power of the selected voxels, obtaining one power value for each condition and each frequency band.

#### 2.1.6 hMMG analysis of emotional facial expressions

Differently from the filtering strategy adopted for the hMMG-EMG correlation analysis, we band-pass filtered the MEG signal from 25 to 150 Hz, being the frequency range in which the EMG power spectrum showed the highest activity and best correlated, on average, with the correspondent MEG sensors (see § 3.1.1 and § 3.1.2).

A the time-frequency analysis of the virtual sensors was computed on the whole band-passed signal (from 25 to 150 Hz, in steps of 5 Hz, ‘mtmconvol’ method option of the ft_frequencyanalysis Fieldtrip function), and on the time-window from −0.5 to 2.5 seconds after the picture presentation (taper = ‘Hanning’, sliding window 0.3 seconds). The result of this procedure was a three-dimensional matrix (virtual sensor x frequency range x time). Data were averaged across frequencies and, the baseline time-window was averaged between −0.5 and 0 seconds, while the target time-window was averaged between 0.5 and 2.5 seconds; this way we obtained, for each participant, two vectors of virtual sensors containing a single power value. In order to avoid any possible artifactual effect due to the estimation of different spatial filters for different conditions, the baseline-time data were used to normalize the data in the time-window of interest (‘db’ baseline from ft_freqbaseline function).

Statistical analyses were performed comparing at a group level each of the “active emotions” (Joy, Fear, Disgust and Sadness) against the neutral one. Then, all combinations involving active emotions were compared with each other (P< 0.05, two-tailed). For presentation purposes we thresholded pictures at t >= |5|.

### 2.2 Passive Observation Task: *Materials and Methods*

#### 2.2.1 Stimuli

We selected 100 pictures from the IAPS dataset. The IAPS database provides normative ratings on three dimensions: valence, dominance and arousal. Based on the valence feature (lowest valence 1.45, highest valence 8.2), they were divided into 5 categories with 20 pictures, respectively: very negative (median valence rating 1.79 +/- median absolute deviation 0.14) negative (2.45 +/- 0.17), neutral (3.76, +/- 0.40), positive (5.18 +-0.58), and very positive (7.31 +-0.40),

#### 2.2.2 Procedure

Each picture was presented on the screen for 6 seconds, followed by a black screen of 3 seconds (Fig. 1B, *Bottom*). Order of pictures was randomized across participants. Participants were asked to pay attention to the pictures and were told that, at the end of the experiment, they would have had to reply to some questions about the stimuli.

#### 2.2.3 Valence-Contraction Correlation

We conducted a more conventional analysis with the scope of highlighting head muscles correlating with picture valences. On the “post-stimulus” dataset we linearly correlated (Pearson) the valence of each picture with the averaged amplitude of the 25-150 Hz frequency window for each virtual sensor, thus obtaining a correlation value for each virtual sensor for each participant.

To test the statistical significance of the correlation values, we used a nonparametric randomization procedure in order to control for multiple comparisons over sensors (Maris & Oostenveld, 2007) implemented in Fieldtrip.

We compared the obtained values against 0 correlation (settings: cfg.method = ‘montecarlo’, cfg.clusterthreshold = ‘nonparametric-individual’, cfg.clusterstatistics = ‘max’, cfg.numrandomization = 5000, 15.5 neighbors on average, method = ‘distance’, neighbor distance = 1.5 cm).

Statistical threshold was set at alpha = 0.05, one-tailed, since we only focused on negative correlations, i.e., the lower the pleasantness of the picture, the higher the muscle contraction. Our results are in agreement with the literature, showing negative linear correlation between regions close to *m. Corrugator supercilii* and hedonic picture ratings (Cacioppo et al., 1986).

#### 2.2.4 Valence classification based on hMMG activity

Data preprocessing followed almost identical steps as the hMMG analysis of emotional facial expressions, with two notable differences: 1) spatial filters were estimated using the whole cohort of trials, i.e., including trials from all conditions. In this case the no baseline correction was necessary, given that the spatial filter applied is identical for all trials. 2) data were epoched from −7 to 7 seconds with respect to picture presentation onset. Data included in the time window −7 to −1 seconds were averaged both along the time and frequency domain (mean frequency = 87.5 Hz) to form the “pre-stimulus” time-window. The same averaging was applied between 1 and 7 seconds after stimulus presentation onset (note that spontaneous facial expression typically exceeds the stimulus offset) to form the “post-stimulus” time-window.

The result of these data processing steps is a two dimensional matrix of virtual channels containing two power values, one from “pre-stimulus” and the other from “post-stimulus” time periods.

A linear discriminant analysis (LDA) was used to classify the data adopting a cross-validation approach provided by the CoSMoMVPA Matlab toolbox (Oosterhof, Connolly, & Haxby, 2016). We conducted a k-fold (k = 10) cross-validation, with 90 pictures used as training-set for the classifier and 10 as test-set. The accuracies resulting from the ten cross-validation procedures were averaged, providing one accuracy value per participant.

Two paired-sample *t*-tests against chance level accuracy (0.2) were performed on the accuracy of the “pre-stimulus” and the “post-stimulus” time windows at group level.

#### 2.2.5 Searchlight Classification

In order to identify areas on the surface of the head carrying most of the information about the picture valence category, a searchlight virtual sensor pattern analysis procedure was carried out on the “post-stimulus” time-window. The searchlight procedure follows similar steps as the previous classification analysis, but with the difference that, instead of considering the whole set of virtual sensors as a features for the classifier, it takes each virtual sensor along with its neighbors (8 on average) as features (called the searchlight), to perform the cross-validation analysis as before and then repeats the same procedure for the next virtual sensor (Kriegeskorte, Goebel, & Bandettini, 2006). Thus, differently from the hMMG analysis of emotional facial expressions, which produced one value of accuracy per participant, the result of the searchlight procedure provides a value of accuracy for each virtual sensor, reflecting the amount of information contained in each virtual sensor and its immediate neighbors.

To statistically evaluate the results of the searchlight classification procedure, we used a threshold-free cluster estimation (Smith & Nichols, 2009) algorithm implemented in CoSMoMVPA (E = 0.5, H = 2). We compared, at group level, the accuracy of each searchlight against the chance value of 0.2.

## 3 Results

The experimental session was divided into two parts, “Passive Observation” and “Imitation” tasks (Fig. 1). While experimentally the two tasks have been run in the above described order, with the aim of not directing participants’ attention to their own facial expressions during the Passive Observation task, we will describe results in the reverse order, i.e., first Imitation followed by Passive Observation task.

### 3.1 Imitation Task

In the Imitation task participants (*N* = 22) were asked to imitate five facial expressions (Joy, Disgust, Fear, Sadness, Neutral) performed by an actress on the screen (Radboud Face Database, RaFD, Langner et al., 2010, Fig. 1A). For comparison purposes with conventional EMG, ten participants wore electrodes in correspondence of left *m. Corrugator Supercilii*, right *m. Frontalis*, and right *m. Zygomaticus Major* (Fig. 2A). The general purpose of the Imitation task was to provide an important quality check that the signal recorded from face muscles was well reconstructed in the hypothesized face locations.

**Fig. 2.**
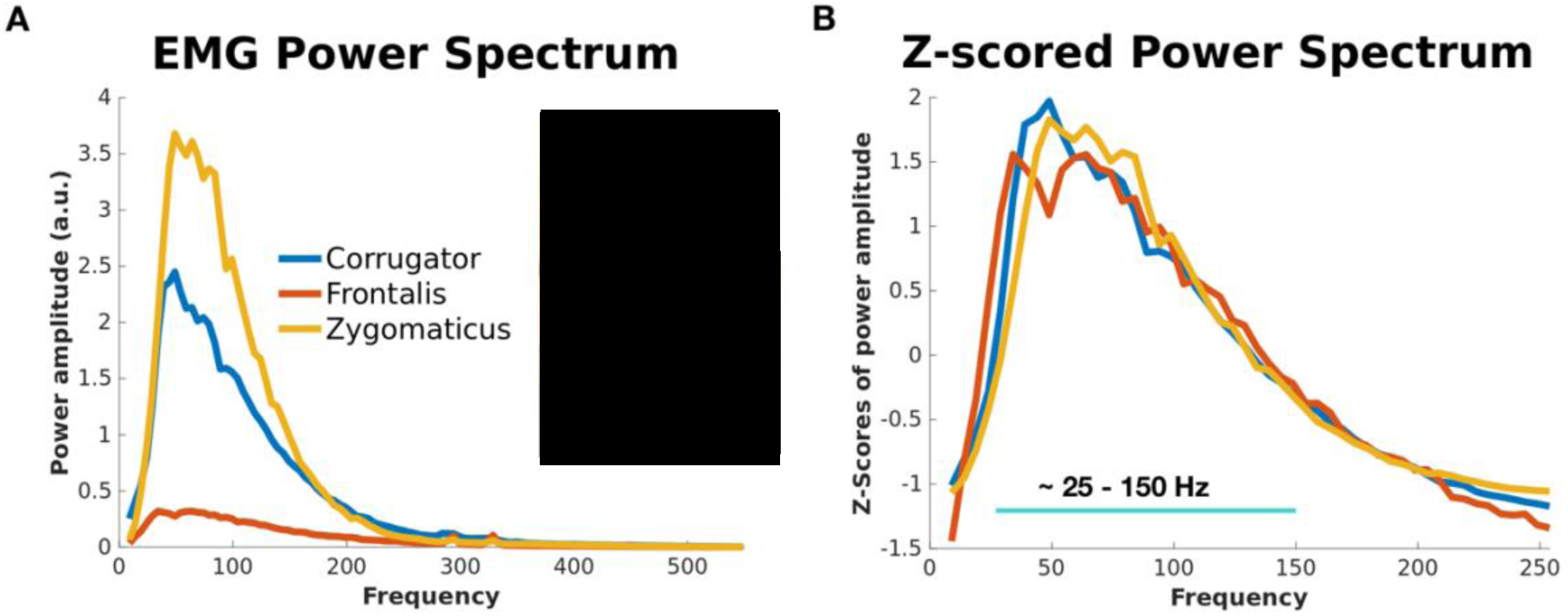
Location and power spectra for the 3 EMG channels. *A):* Average power spectrum across participants of the 3 EMG channels representing the power during the imitation conditions that are best associated with their contraction: left *m. Corrugator Supercilii* during Disgust imitation, *m. Frontalis* during Fear imitation, *m. Zygomaticus Major* during Joy imitation. *B):* Z-score transformation of individual EMG power spectra shows that the channels have very similar spectral components. The band containing dominant frequencies, between 25 and 150 Hz, has been used in the following analyses. <u>*images have been removed/obscured due to a bioRxiv policy on the inclusion of faces</u>

#### 3.1.1 EMG Power Analysis

Being the gold-standard in muscular signal recording, as a first step, a spectral characterization of the muscular activity picked up by EMG was performed, in order to inform the subsequent MEG source localization analysis. The resulting power-spectrum shows strongest signal in the 25-150 Hz range, peaking between 35 and 45 Hz (Fig. 2A). The highest overall power is reached by *Zygomaticus Major* and *Corrugator Supercilii*, followed by *Frontalis* muscles, which show the lowest absolute power. However following a Z-score transformation of the power spectra of each muscle, the channels show virtually identical spectral power distribution. Based on the z-scored EMG power spectra, we selected a frequency band ranging between 25 and 150 Hz to be used in all the following analysis.

#### 3.1.2 hMMG-EMG Correlation

If hMMG is to be used as a tool to record muscular activity localized on the face, we would expect a strong correlation at relevant locations with signals recorded using conventional EMG. To test this we conducted a correlation analysis between band-passed filtered hMMG and EMG time-courses; an example of such band-passed time-series is provided in figure 3A. EMG signal from *m. Corrugator Supercilii and* hMMG signal were correlated during imitation of Disgust, while EMG from *m. Frontalis*, and *Zygomaticus Major,* respectively, were correlated with hMMG during Fear and Joy imitation. We selected the 2.5% (approximately 118 out of 4735 voxels) of the highest correlation values for each participant, in each condition and frequency band. Figure 3B shows the number of counts representing the number of participants having a virtual sensor among the highest 2.5% in correlation values for different frequency bands. The results show that overall the location of the highest count number matches the “a-priori” EMG electrode placement, being in close proximity to the position of the targeted muscles.

**Fig. 3.**
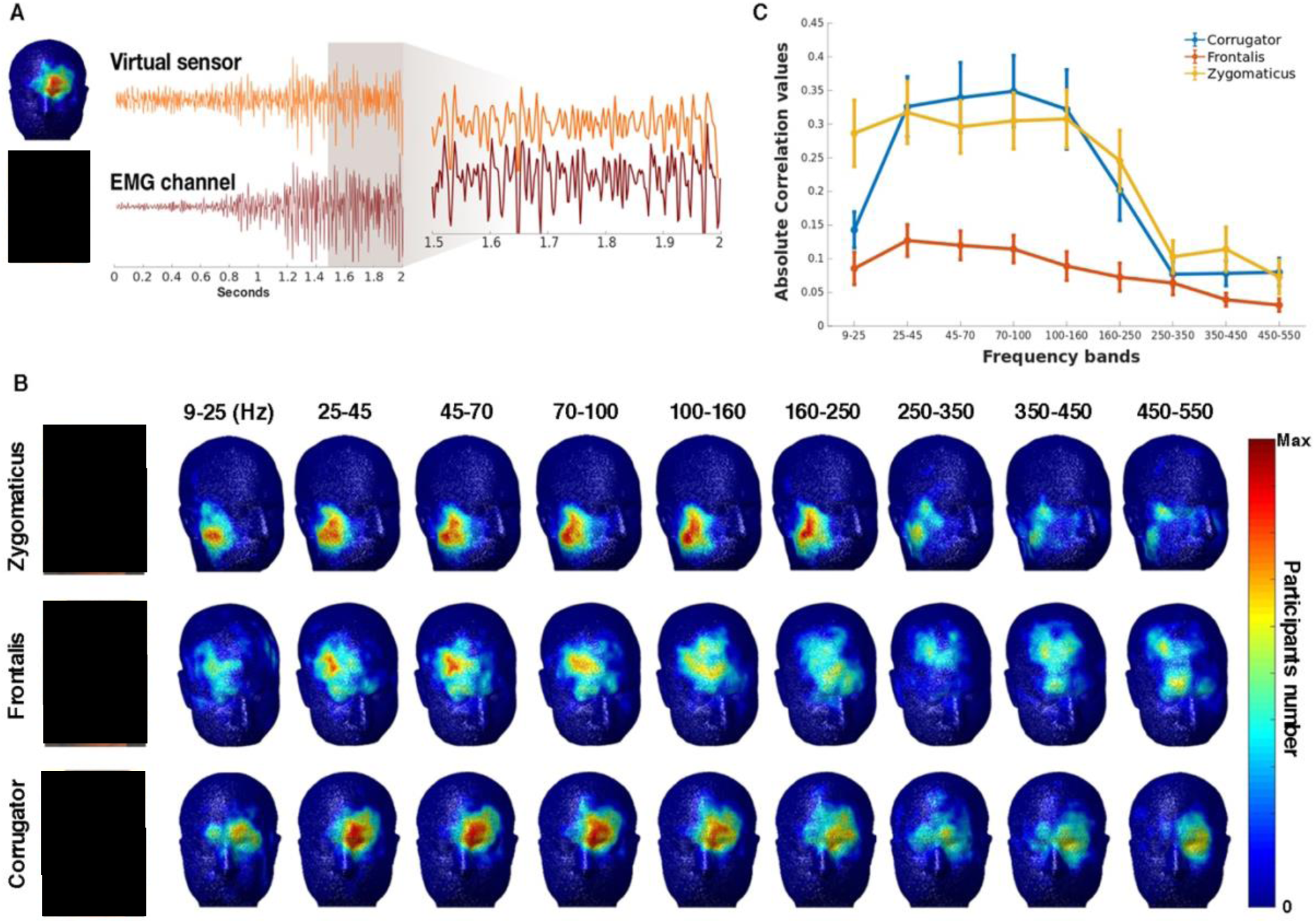
A) Example trial hMMG vs. EMG time-course. The two traces represent EMG (*m. Corrugator Supercilii*) and a correspondent hMMG virtual sensor during a single contraction; magnification shows the similarity of the two traces during the time-window from 1.5 and 2 second after picture presentation. MEG data have been band-pass filtered in the 25-150 Hz range before being submitted to LCMV beamformer. B) **hMMG-EMG localization**. hMMG-EMG absolute correlation values have been computed separately for each participant wearing EMG electrodes. The upper row shows the number of times in which a virtual sensor was among the 2.5% with the highest correlation during Disgust imitation. The middle and the bottom rows show the same metric during Fear and Joy imitation. Correlations were computed after time-series had been band-pass filtered in 9 different frequency bands. C) **hMMG-EMG average correlation**. The absolute correlation values have been averaged from virtual sensors containing the highest 2.5% absolute correlation values for the highest number of participants down to the highest number – 1, separately for each condition, after a 25-150 Hz band pass filtering (e.g., if maximum count in the correlation after 25-150 Hz band pass filtering for one condition was 9, then all the channels having 8 or 9 counts were selected). After the relevant voxels were obtained, means and standard error of the correlation per each frequency band were calculated. <u>*images have been removed/obscured due to a bioRxiv policy on the inclusion of faces</u>

After having localized the voxels which consistently showing the highest correlations with the EMG signal, we wanted to measure in absolute terms the average magnitude of the correlations. Figure 3C shows the distribution of the average hMMG-EMG correlation values across frequency bands by considering virtual sensors showing the highest count and highest count – 1, separately for each muscle (e.g., if maximum count for one condition was 9, then all the channels having 8 or 9 counts were selected, and their correlation values averaged). For all channels recorded, the highest correlation values laid approximately between 25 and 150 Hz, in line with the EMG power analysis results. Within that frequency range, *Zygomaticus* and *Corrugator* muscles had the highest correlation values, between 0.296 and 0.317 and between 0.322 and 0.349, respectively. The *Frontalis* muscle has overall smaller correlation values compared to the other two muscles, ranging between 0.127 and 0.09 in the 25-150 Hz band. Overall, the results of this analysis established a strong similarity between hMMG based muscular recordings and standard EMG.

#### 3.1.3 hMMG analysis of emotional facial expressions

To illustrate the potential of our approach to measure facial muscular activity, we performed statistical contrasts of the hMMG signals in order to obtain topographic differences between emotional facial expressions. In a first step we contrasted emotional expressions to the neutral expression in order to test whether commonly reported muscle groups can be identified. As figure 4A shows, the expected muscular activity characterizing each emotion was detected; Joy imitation shows peaks at inferior-lateral locations on the face, in proximity to *m. Zygomaticus Minor, m. Zygomaticus Major* and *m. Risorius,* while Disgust and Fear imitation showed peaks in correspondence to lower (*m. Corrugator Supercilii, m. Procerus*) and higher forehead (*m. Frontalis*), respectively. Notably, other portions of the head, outside those predicted, were activated for Joy and Fear imitation; areas close to the ears show a peak of activity in Joy imitation, possibly pointing to *Anterior Auricularis* muscles contractions, while in Fear imitation activity in correspondence to neck muscles was detected. Comparisons between “active emotions” (Fig. 4B) are in line with observations shown in figure 4A and highlight specific areas of muscular activation differences between emotion expressions.

**Fig. 4.**
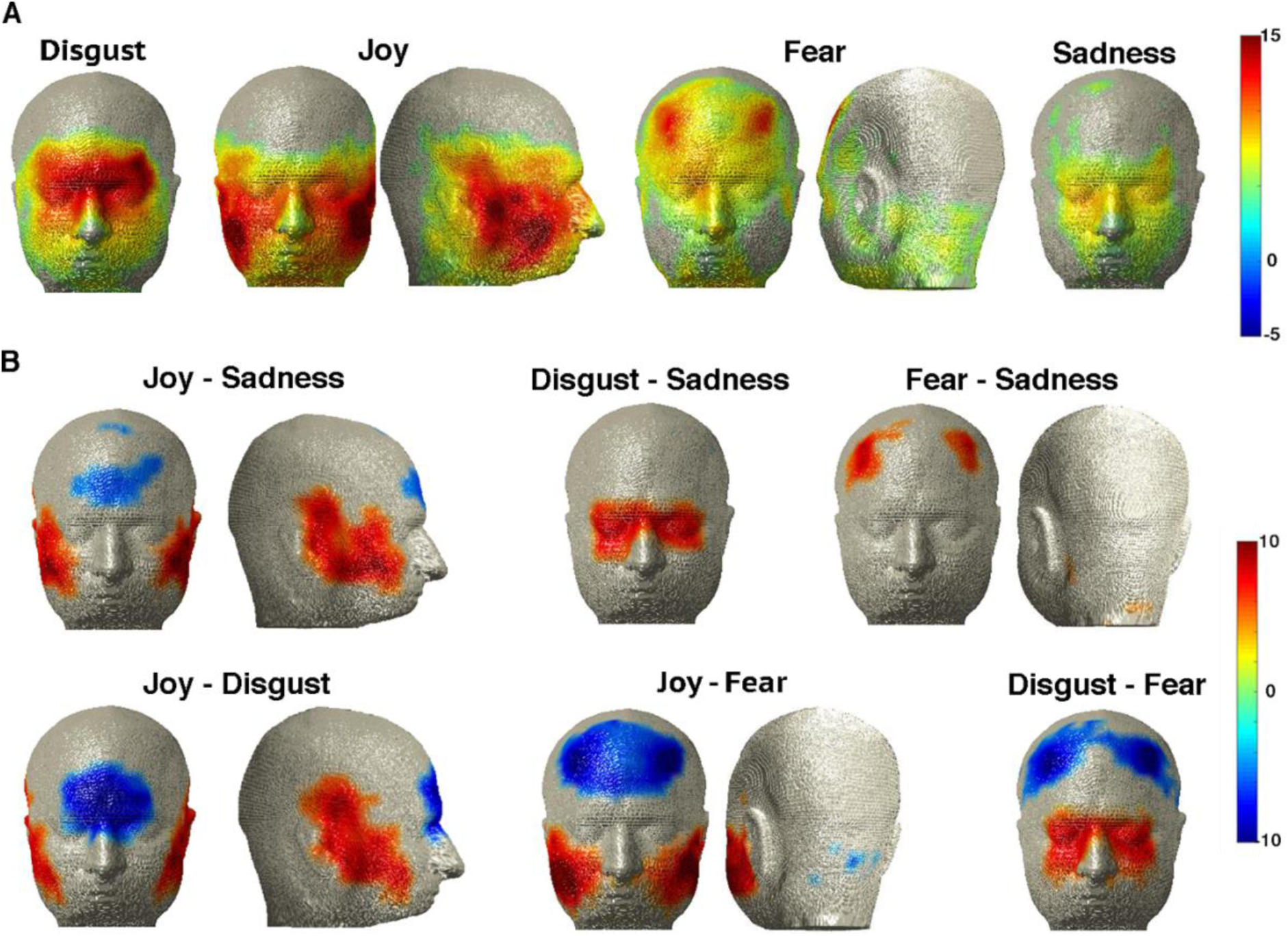
*(A)* hMMG of the comparisons emotion vs. neutral expression. *(B)* **hMMG of the comparisons of emotion expressions** against each other. Virtual sensors showing *t*-values above |5| are displayed.

These results prove that hMMG is suited for localizing muscular activity during facial expressions, but most importantly, they highlight the advantage of “whole-head” recordings against the classical “a-priori” electrode selection, revealing that unexpected or overlooked muscles contributed to the expression patterns.

### 3.2 Passive Observation Task

During imitation, muscular activity is particularly intense, so the usefulness of hMMG in a setting with more subtle muscle activity still needs to be established. This was the purpose of the Passive Observation Task, where participants (N = 17) were asked to attentively observe a series of emotion-inducing pictures selected from the International Affective Picture System (IAPS) database ranging from very negative to very positive valence ratings (Bradley & Lang, 2007).

We conducted a “Valence-Contraction Correlation” analysis, in which, for each participant the amount of activity of each virtual sensor was correlated with the normative valence rating of the pictures presented (Fig. 5A), followed by a classification analysis based on hMMG activity (Fig. 5B), and a “Searchlight classification” (Fig. 5C).

**Fig. 5.**
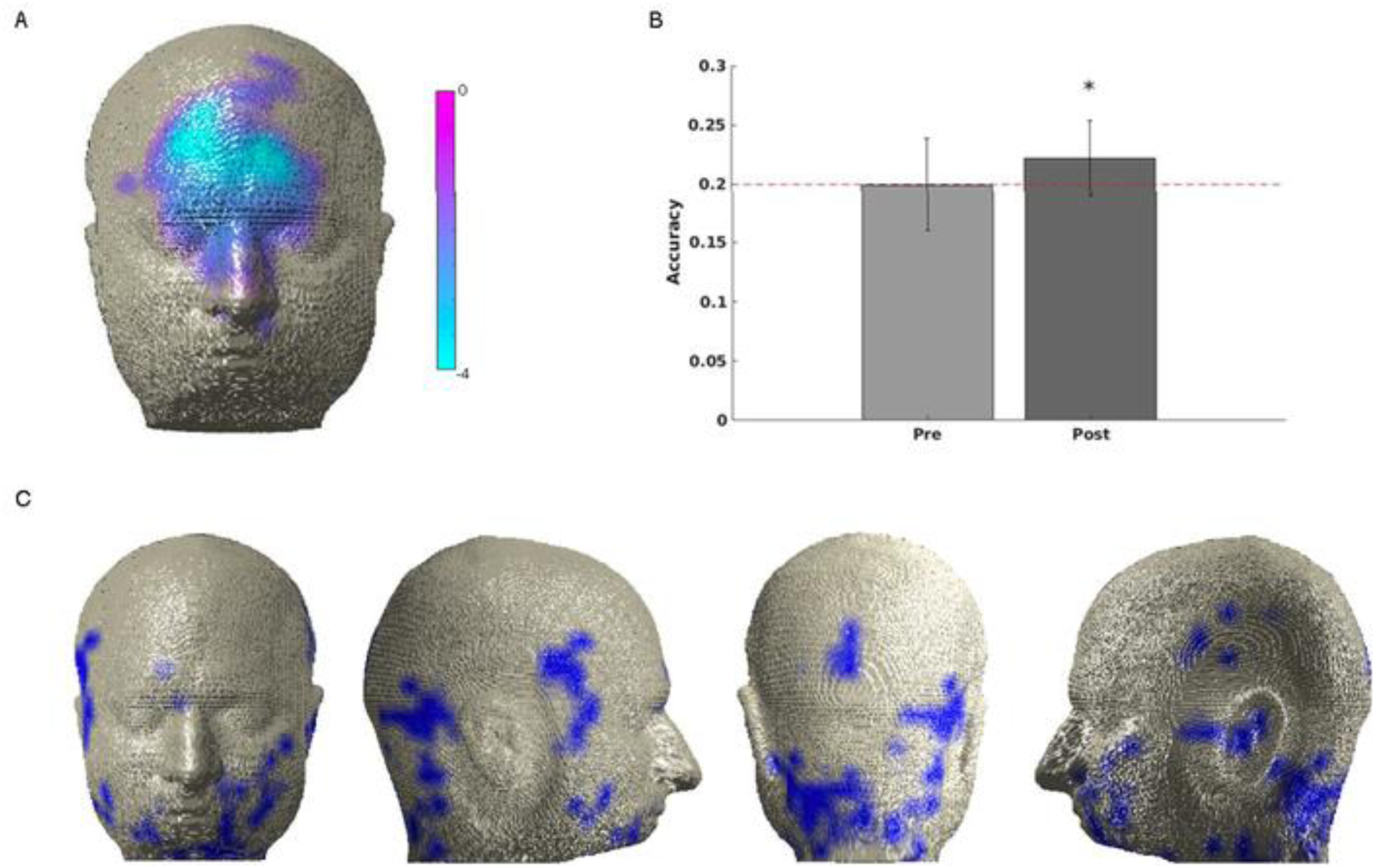
A) Valence – Contraction correlation: the only statistically significant cluster was found in the region including bilaterally the *Corrugator* muscles, but also regions higher on the z-axis, a picture resembling the Joy–Disgust comparisons but also including areas in proximity to the medialis pars of frontalis muscles. The lateral bar represents the values of the *t*-test against 0 correlation. B) **Accuracy values on the whole-head voxel:** average classification accuracies in the pre-stimulus and the post-stimulus period. Error bars represent standard error of the mean. C) **Searchlight classification results**: A hub of information is provided within the neck region. Only voxels showing P<0.005, are represented.

#### 3.2.1 Valence-Contraction Correlation

To validate hMMG, it is important to illustrate that it is sensitive to established effects. To this end we aimed to replicate a well-established result (Cacioppo et al., 1986; LANG et al., 1993; Larsen et al., 2003) from studies using conventional EMG, showing that amplitude of *m. Corrugator Supercilii* contractions is negatively correlated with the valence of the perceived stimulus. Our cluster-based statistical procedure confirmed this finding, yielding one negative cluster of voxels (P<0.05 one-tailed) located medially on the upper face, including *m. Nasalis, Corrugator Supercilii* bilaterally, and *m. Frontalis pars medialis* bilaterally (Fig. 5C), confirming the similarity between the classical EMG results and hMMG recordings.

#### 3.2.2 Valence classification based on hMMG activity

Classical data from the literature, as the valence-contraction correlation, extracted information from the channels one at a time. hMMG instead, comprising multiple voxels, is suited to take advantage of classifiers to characterize informative multidimensional patterns of activity. To test if hMMG activity patterns could predict the valence of the observed pictures, we sorted the images dividing them into 5 valence classes and applied a Linear Discriminant Analysis (LDA) classifier. The analysis yielded, for the post-stimulus time window, an accuracy level above chance level (0.2), indicating that valence can be classified significantly above chance (0.222; *t*(16) = 2.70, *P* = 0.016, Fig. 5B). For the pre-stimulus period accuracy no significant difference from chance was detected (0.199; *t*(16) = −0.05, *P* = 0.962). Since the previous analysis used all grid points simultaneously to classify valence category, no spatial information was obtained. For this purpose we used a searchlight approach, yielding a spatial distribution of accuracy. This analysis showed that valence categories are distinguishable above chance in different regions of the head, including lower and lateral portion of the face, and notably, neck regions (Fig. 5C).

## 4 Discussion

Given its ease of application, usefulness, and low cost, EMG is the gold-standard of muscular recording. As any other technique however, EMG suffers from limitations; here we focused especially on one of them, namely, the need for “a-priori” selection of the electrodes to be placed on the participant’s face. In order to overcome this limitation, we used conventional whole-head MEG as a large-scale EMG recorder, providing magnetomyographic recordings of the whole head at once.

### 4.1 hMMG provides similar information as EMG with high spatial coverage

As a first step, to test the hypothesis that hMMG is capable of measuring and locating muscular activity at circumscribed regions, we calculated the correlation between the EMG and hMMG for facial expressions of Joy, Disgust and Fear. Clearly, if the MEG source space reconstruction of head muscular activity was spatially unspecific, then a dispersed pattern of hMMG-EMG correlation would have emerged with no consistent peak locations. Instead, we were able to find clear peaks of correlations among participants, located in correspondence to the muscles we expected to be correlated with the EMG activity. Secondly, we showed that the correlation between EMG and hMMG time-courses was modulated by the frequency range of the initial band-pass filtering of the traces, and that correlation followed a trend, across filtering frequency bands, that resembled the one found on the EMG power spectrum. Indeed the highest correlation values were found within the band corresponding to the 25-to-150 Hz frequency range, analogous to the pattern found in the power spectrum of the EMG signal. If the two signals were unrelated, no pattern would have been found across frequency bands. Instead, our results clearly point to a relationship between EMG and hMMG signals. Thirdly, we showed that the patterns of muscular activity detected by the hMMG across the imitation task are highly compatible with expected facial expression patterns, with the addition of unpredicted activations, as shown for the case of *m. Auricularis* and neck muscles, an outcome which highlights the advantage of having a large-scale muscle recorder compared to an “a-priori” and limited channel selection.

Overall these findings illustrate that the hMMG can be used to reconstruct muscular information at circumscribed locations from the entire head. The latter aspect is the central added value of hMMG over or in conjunction to conventional EMG. Our results also indicate muscular activity in head regions that are challenging to detect using classical bipolar EMG (e.g., due to presence of hair) such as *Auricolaris* muscles in proximity of the ears, or *Occipitalis* muscles on the back of the head.

### 4.2 hMMG detects subtle muscular activity patterns during affective picture viewing

While the application of hMMG might potentially expand beyond basic research, our study already shows the benefit for affective neuroscience research. In order to show consistency between hMMG and conventional EMG results in this domain, we correlated the activity at each voxel with the valence of the pictures observed, aiming to reproduce the well-known result of a negative linear correlation between valence and *m. Corrugator Supercilii* activity (Cacioppo et al., 1986; LANG et al., 1993; Larsen et al., 2003). A significant negative correlation was observed comprising *m. Corrugator Supercilii* bilaterally, extending to upper fibers, possibly in the *pars medialis* of the *Frontalis* muscle. Considering the Facial Action Coding System as an atlas for facial muscles contraction description, this pattern resembles a mix of activity in action-unit 1 and 4 (inner brow raiser and brow lowerer Ekman & Friesen, 1978) suggesting that the view of negative-valence pictures went along with sadder emotional states. To exploit the advantage of having multiple voxels recorded at one time, we applied an LDA to the hMMG data aiming to predict the valence category of the presented emotion-inducing pictures. Using whole-head patterns enabled the above chance classification of a presented image’s valence category. As a last step, the use of hMMG in combination with a searchlight analysis uncovered informative muscle activity patterns in the neck region, previously overlooked when studying the relationship between emotional experience and muscle contraction. It is worth noting that, differently from EMG, no electrodes have to be attached to the participants’ face when using hMMG; this characteristic avoids the interaction between electrodes and skin, which might alter proprioceptive feedback and thus influencing emotional facial expression. In addition, it might draw participants’ attention to their face and thus reduce spontaneity of facial expression. Overall, this set of promising results show how hMMG can be readily utilized as neuroscientific tool for affective science, to significantly exceed the amount of information that can be acquired using conventional EMG alone.

### 4.3 Limitations and Caveats

The correlation analysis between EMG and hMMG provided intermediate values, with the highest average correlation of around 0.3. This value reached up to 0.7 for some participants, but overall it is clear that the hMMG does not model the EMG time series perfectly. Several reasons for this discrepancy exist: 1) EMG spatial selectivity might have been influenced by cross-talk, the phenomenon by which the EMG signal, aimed to record only one muscle, is contaminated by the activity of surrounding muscles. In hMMG, muscular activity is estimated at exact point sources using an adaptive spatial filter. 2) The signal-to-noise ratio is higher for the EMG compared to the virtual sensor traces as suggested by the pre-contraction period in Fig. 3A. Some contributing reasons could be vicinity of EMG electrodes to the source or the measurement via a bipolar montage. 3) We used a template MRI for almost half of the participants, which could have reduced average signal-to-noise ratio for hMMG. 4) While we could have chosen to correlate the envelopes of hMMG and EMG time-series, we opted for the stricter comparison, by directly correlating the raw time-series. 5) Performing emotional expressions was not easy for many participants, since they were only briefly trained to perform those expressions before the experiment started; this problem might have resulted in non-optimal contractions throughout the imitation, leading to the production of poor muscular signal, which might have reduced correlation values.

Related to the last point, it is obvious that some expressions were performed better than others: the Sadness map does not correspond entirely to the pattern of activity hypothesized a-priori, which would have been characterized by *m. Frontalis* (pars-medialis), plus *m. Corrugator Supercilii* activity. To a weaker extent this also applies to Fear expressions. While, as a first thought, this lack of correspondence might be attributed to intrinsic limitations of the hMMG technique, the most likely explanation can be found in the participants difficulty in imitating emotional expressions. As a consequence, this difficulty should not only be reflected in the hMMG but also in the EMG signal; indeed, for example, the average EMG activity recorded on the Frontalis muscle in Fear expressions was almost five times lower than the one recorded from the *m. Corrugator*, so clearly differences like these might emerge also in the hMMG signal.

It is also worth mentioning that source space activity maps for the imitation task as well as statistical maps have been created based on only 35 samples per emotion, which is a small number compared to classical experimental MEG-brain designs.

Altogether, all these considerations suggest that conventional EMG is advised whenever the relevant muscle is precisely known in advance and confounding due to proprioceptive feedback and emotion self-awareness is not of concern. In all other cases, hMMG is a powerful alternative or addition. hMMG is just at the beginning of its development, which leaves space for important technical improvements that might provide a more fine-tuned identification of muscular activity patterns and signal-to-noise ratio. For example, we will exploit a-priori information from muscular atlases and the orientation of the magnetic dipoles produced by muscles to better separate close muscle fibers.

### 4.4 Conclusions and Future Perspectives

This study demonstrates that hMMG is a powerful method to monitor whole-head muscular activity, yielding some advantages over classical EMG. In addition, our data show the potential of hMMG in psychological experiments, by replicating and extending established findings in emotion research; in this context hMMG can be readily used as a stand-alone muscular recording technique. Beyond the illustrated possibilities, hMMG holds promise as a clinical screening tool; e.g., using hMMG as an adjunct to electroneurography might provide a deeper characterization of facial palsies, identifying more easily the muscular groups affected by such pathologies (electrical stimulation inside the MEG has been shown to be feasible by our group (Neuling et al., 2015). hMMG might also become an alternative to monitoring the innervation process of implanted muscle flaps after “facial animation” procedures (Bianchi, Copelli, Ferrari, Ferri, & Sesenna, 2010); normally, this monitoring is performed by implanting a needle for EMG recording inside the innervated flap, which is uncomfortable especially for children. Another advantage over EMG, which in a clinical context might become critical, is that hMMG might record activity of muscles that are difficult to reach with standard bipolar surface or even needle recordings (e.g., *Stapedius* muscle inside the inner ear, the extraocular muscles, muscles involved in swallowing or located inside the larynx, and inner neck muscles such as *m. Longus Colli*). However, given the inexpensive and easy application of EMG, in many cases, hMMG may be assisted by EMG, combining the strengths of both approaches. Overall, we believe that hMMG is likely to gain prominence as a muscular recording technique, expanding its advantages not only to the affective sciences domain, but also to applied contexts such as the clinical one.

## Acknowledgments

The first author was supported by Lise Meitner FWF grant M 2171-BBL.

We would like to thank Manfred Seifter and Viola Heberger for their help in collecting the data

